# One-step assembly of large CRISPR arrays enables multi-functional targeting and reveals constraints on array design

**DOI:** 10.1101/312421

**Authors:** Chunyu Liao, Fani Ttofali, Rebecca A. Slotkowski, Steven R. Denny, Taylor D. Cecil, Ryan T. Leenay, Albert J. Keung, Chase L. Beisel

## Abstract

CRISPR-Cas systems inherently multiplex through their CRISPR arrays--whether to confer immunity against multiple invaders or by mediating multi-target editing, regulation, imaging, and sensing. However, arrays remain difficult to generate due to their reoccurring repeat sequences. Here, we report an efficient, one-step scheme called CRATES to construct large CRISPR arrays through defined assembly junctions within the trimmed portion of array spacers. We show that the constructed arrays function with the single-effector nucleases Cas9, Cas12a, and Cas13a for multiplexed DNA/RNA cleavage and gene regulation in cell-free systems, bacteria, and yeast. We also applied CRATES to assemble composite arrays utilized by multiple Cas nucleases, where these arrays enhanced DNA targeting specificity by blocking off-target sites. Finally, array characterization revealed context-dependent loss of spacer activity and processing of unintended guide RNAs derived from Cas12a terminal repeats. CRATES thus can facilitate diverse applications requiring CRISPR multiplexing and help elucidate critical factors influencing array function.

## INTRODUCTION

CRISPR-Cas systems represent RNA-directed immune systems whose programmable nucleases have become powerful technologies for genome editing, gene regulation, imaging, and diagnostics (Barrangou and Doudna, 2016; Komor et al., 2017). CRISPR technologies derive from an increasing assortment of CRISPR-associated (Cas) single-effector nucleases (Koonin et al., 2017; Mohanraju et al., 2016). These nucleases include the originally discovered and widely-used Type II Cas9 nucleases that introduce a blunt cut into dsDNA targets (Gasiunas et al., 2012; Jinek et al., 2012), the more recently discovered Type V Cas12a (or Cpf1) nucleases that introduce a staggered cut into dsDNA targets and degrade ssDNA upon target recognition (Chen et al., 2018; Zetsche et al., 2015), and the functionally unique Type VI Cas13a (or C2c2) nucleases that cut ssRNA targets and can non-specifically degrade cellular RNAs upon target recognition (Abudayyeh et al., 2016; Shmakov et al., 2015). Across this diversity, CRISPR-Cas systems share an inherent capacity for multiplexing through their CRISPR arrays. These arrays are composed of alternating conserved “repeats” and targeting “spacers”, where some prokaryotes can encode up to a few hundred spacers in a single array that are stable over evolutionary timescales (Jansen et al., 2002). The transcribed array undergoes processing into multiple guide RNAs (gRNAs) derived from each repeat-spacer pair. Each gRNA directs the Cas nuclease to bind and cleave complementary DNA or RNA targets flanked by a short protospacer-adjacent motif (PAM) or a protospacer-flanking sequence (PFS) (Leenay and Beisel, 2017).

The multiplexing capacity of CRISPR arrays was well recognized before the advent of CRISPR-Cas9 technologies (Barrangou et al., 2007; Brouns et al., 2008). However, multiplexing with Cas9 outside of bacterial systems was constrained by the limited portability of RNase III and the trans-activating CRISPR RNA (tracrRNA) necessary for processing of Cas9 arrays (Cong et al., 2013). The invention of the single-guide RNA (sgRNA) obviated the need to co-express the tracrRNA and RNase III (Jinek et al., 2012), although the sgRNA sacrificed the inherent multiplexing capability of CRISPR arrays and therefore required numerous engineering workarounds to produce multiple sgRNAs (Nissim et al., 2014; Wong et al., 2016; Xie et al., 2015). The discovery that the Cas12 and Cas13 nucleases processed CRISPR arrays through their own endonucleolytic domains (East-Seletsky et al., 2016; Fonfara et al., 2016; Zetsche et al., 2015; Zhong et al., 2017) intensified the pursuit of synthetic CRISPR arrays for multiplexing applications. More recent examples include multiplexed genome editing and gene activation with Cas12a (Tak et al., 2017; Zetsche et al., 2016), and multiplexed gene silencing with Cas13a (Abudayyeh et al., 2017). Beyond these recent demonstrations, use of arrays could also advance many other CRISPR technologies that have yet to employ CRISPR arrays for multiplexing, such as paired nickases or FokI fusions (Guilinger et al., 2014; Tsai et al., 2014), enhanced gene drives (Noble et al., 2017), proximal CRISPR targeting (Chen et al., 2017a), multi-pathogen antimicrobials (Fagen et al., 2017), multiplexed base editing (Banno et al., 2018), combinatorial nucleic-acid sensing with Cas12a and Cas13a (Chen et al., 2018; Gootenberg et al., 2017), and combinatorial screens (Billon et al., 2017; Kuscu et al., 2017; Peters et al., 2016). However, one of the major barriers impeding multiplexing applications is how to generate CRISPR arrays.

As a default, CRISPR arrays would be chemically synthesized as linear dsDNA by commercial vendors. Unfortunately, the reoccuring repeat sequences inherent to these arrays currently pose major technical complications when assembling individually synthesized oligonucleotides, resulting in vendors regularly rejecting customer requests even for a minimal single-spacer array. Gene synthesis has offered a more reliable means of obtaining custom CRISPR arrays. However, synthesis often comes at large cost (~5x the price of a linear dsDNA) and timeframes (~1 month), and the synthesis can often fail. As an alternative, a few groups have developed different assembly methods based on annealing shorter oligonucleotides into repeat-spacer subunits that can be assembled sequentially or simultaneously into arrays (Table S1) (Cress et al., 2015; Gomaa et al., 2014; Tak et al., 2017; Vercoe et al., 2013; Zetsche et al., 2016). For instance, one study sequentially inserted individual repeat-spacer subunits into a non-target spacer to generate Cas9 arrays with up to a three spacers (Cress et al., 2015) while another assembled Cas12a arrays with up to three spacers in one step by creating 5’ overhangs that fall within different parts of the conserved repeat (Tak et al., 2017). While these approaches were used to successfully generate CRISPR arrays harboring 2 - 4 spacers, they cannot scale to larger arrays and often exhibited low cloning efficiencies even for these small arrays (Table S1). Therefore, the ability to easily, cheaply, and quickly generate CRISPR arrays remains an impediment to the widespread use of CRISPR multiplexing and the fundamental study of array processing and function.

Here, we present an assembly scheme for the efficient, one-step generation of large CRISPR arrays. The method, which we have named CRATES (<u>CR</u>ISPR <u>A</u>ssembly through <u>T</u>rimmed <u>E</u>nds of <u>S</u>pacers), relies on ligating ~60-nt repeat-spacer units at defined junctions within the trimmed and therefore expendable portion of each spacer. The junctions allowed for the efficient assembly of arrays with up to seven spacers. Using the resulting arrays, we demonstrated multiplexed nucleic-acid cleavage and gene regulation with the single-effector nucleases Cas9, Cas12a, and Cas13a in the bacterium *E. coli*, the eukaryote *Saccharomyces cerevisiae*, and a cell-free transcription-translation (TXTL) system. Moreover, we created composite arrays that were utilized by more than one nuclease, where these arrays were used to demonstrate enhanced targeting specificity through the coordinated cleavage of an on-target site by Cas9 and blocking of multiple off-target sites by a catalytically-dead Cas12a. Finally, interrogation of the arrays revealed design considerations and challenges, such as widely ranging abundances of the processed gRNAs that could render an otherwise functional spacer inactive and Cas12a deriving unintended gRNAs from the terminal repeat. In total, the assembly scheme is expected to streamline multiplexing with numerous CRISPR single-effector nucleases and facilitate the interrogation of the processing, function, and evolution of CRISPR arrays across the prokaryotic world.

## RESULTS

### CRATES: a one-step assembly scheme for Class 2 CRISPR arrays

Given the growing interest in multiplexing with Cas12a for genome editing and gene regulation, we first sought to develop a one-step assembly scheme for these arrays that relied on short oligonucleotides and could scale to large arrays. Modular assembly techniques have proven effective for the one-pot assembly of multiple DNA fragments (Casini et al., 2015). These techniques involve the digestion of DNA fragments with Type IIS restriction enzymes, resulting in 4-nt overhangs that are annealed to form the junctions between fragments in the final construct. While the overhangs theoretically can be any sequence, defined sets of overhang sequences are commonly used to prevent the formation of unintended junctions (e.g. between two overhangs that can partially anneal) (Ng and Sarkar, 2012). The question was where to insert these junctions within a Cas12a array without disrupting array processing or CRISPR function. The original characterization of Cas12a revealed that each spacer was trimmed at its 3’ end from 30 nts in the transcribed array to ~23 nts in the processed gRNA (**Fig. 1A**) (Zetsche et al., 2015). Furthermore, recent crystal structures of the Cas12a:gRNA ribonucleoprotein complex bound to target DNA showed that only the first 20 nts of the guide portion of the gRNA participated in base pairing (Swarts et al., 2017; Yamano et al., 2017). We therefore chose the trimmed region of the spacer as the site of the defined junction, because spacers can accommodate virtually any sequence and the junction would not be expected to participate in target recognition.

**Figure 1.**
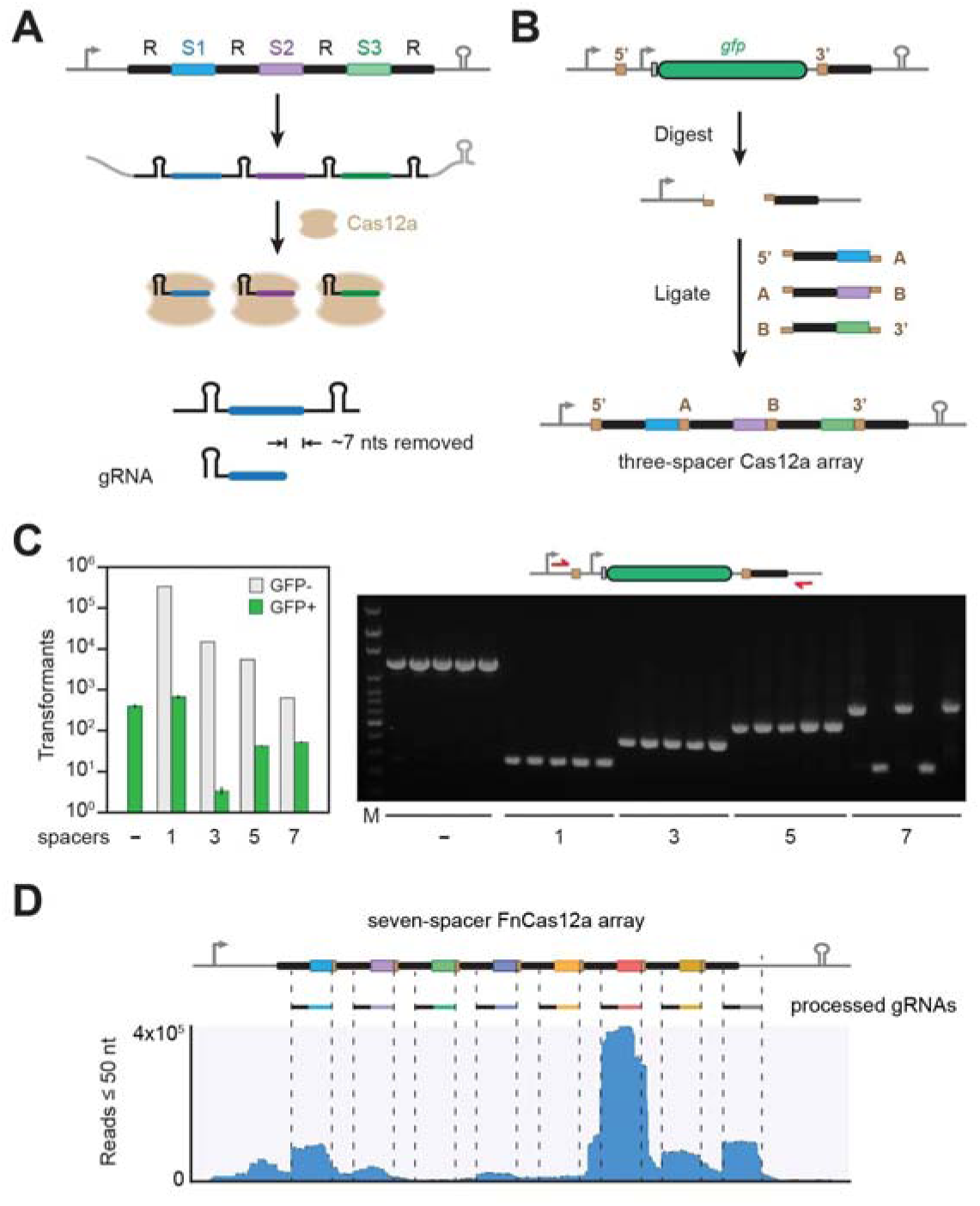
A one-step assembly scheme for CRISPR-Cas12a arrays based on spacer trimming. (A) CRISPR array processing by the Cas12a nuclease. Each array comprises alternating repeats (R, black bars) and spacers (S1-S3, colored bars). Processing includes 3’ trimming of the spacer to form the final gRNA. (B) Cloning scheme for the one-step assembly of multispacer arrays recognized by Cas12a. A GFP-dropout construct is flanked by two Type IIS restriction sites and a 3’ direct repeat. The digested construct is ligated with annealed oligonucleotides encoding individual repeat-spacers in one step. The initial repeat is located at the 3’ end so the resulting array begins and ends with a repeat. The sequence and 5’ or 3’ directionality of the junction overhangs determine the order of assembly. The resulting assembly junctions fall within the trimmed portion of the processed guide RNA and therefore would not be involved in target recognition. (C) Efficient assembly of up to a seven-spacer array. Assembly efficiency was based on the relative proportion of fluorescent or non-fluorescent *E. coli* colonies (left) and the correct insert size of non-fluorescent colonies subjected to colony PCR (right). Values represent the geometric average and S.E.M. from three independent transformations starting from separate colonies. See Table S3 for the specific sequences used. - is the original GFP dropout construct. (D) RNA-seq analysis of the transcribed 7-spacer array. The assembled array and FnCas12a were expressed in an all-E. *coli* cell-free transcription-translation reactions. Reads of no more than 50 nts were mapped to the original array expression construct. Related to Figure S1 and Tables S1-S3.

We next created a base construct for the assembly and expression of Cas12a arrays in the bacterium *Escherichia coli* (**Fig. 1B**). The construct contained a total of six components: a constitutive promoter to drive transcription of the array, two Type IIS restriction sites for inserting multiple repeat-spacer subunits, a GFP reporter construct that is excised as part of array assembly, a 3’ repeat so the final array begins and ends with repeats, and a terminator to halt transcription. The repeat-spacer subunits were formed by annealing two oligonucleotides of ~66 nts, thereby allowing us to readily generate 5’ or 3’ overhangs of any sequence. By alternating 5’ and 3’ overhangs and choosing sets of compatible junctions previously validated for efficient modular assembly (Ng and Sarkar, 2012), we could greatly reduce the frequency of array misassembly. We chose 4-nt overhangs given their use by many modular assembly techniques, although other overhang lengths could be used. The repeat-spacer subunits and the base construct could then be assembled in a one-pot reaction that cycles between digestion of the backbone with a Type IIS restriction enzyme and ligation by T4 DNA ligase. Figure 1B depicts the assembly of a three-spacer array with four distinct junctions (5’, A, B, 3’). We term the resulting assembly method CRATES for <u>CR</u>ISPR <u>A</u>ssembly using <u>T</u>rimmed <u>E</u>nds of <u>S</u>pacers.

### CRATES enabled efficient array assembly and revealed constraints on gRNA production

We first explored how well CRATES could assembly arrays containing different numbers of spacers. We used the 36-nt repeats for the Cas12a nuclease from *Francisella novicida* U112 (FnCas12a), one of the best-characterized Cas12a nucleases to-date (Zetsche et al., 2015), and we designed up to seven spacers containing distinct 26-nt sequences and 4-nt junctions. Following the assembly reaction and transformation into *E. coli*, we counted the relative number of fluorescent or non-fluorescent cells, where non-fluorescent cells lost the GFP expression construct and therefore would be expected to contain assembled arrays (**Fig. 1C**). We found that the total number of transformants decreased for arrays with more spacers, although non-fluorescent colonies always outnumbered fluorescent colonies by a factor of 30 up to 1,000. In contrast, only fluorescent colonies were observed in the absence of any added repeat-spacer subunits. We further performed colony PCR to assess the extent to which non-fluorescent colonies harbored the expected band size (**Fig. 1C**). Testing five random, non-fluorescent colonies yielded the correct band size 100% of the time (5/5) for arrays with up to five spacers and 60% of the time (3/5) for arrays with seven spacers. Sanger sequencing of individual colonies confirmed that the arrays contained the expected sequence. CRATES therefore represents a simple and efficient way to assemble large CRISPR arrays up to and potentially exceeding seven spacers.

We next asked how the seven-spacer array--the largest array we assembled--undergoes processing by FnCas12a when transcribed. We used a recently-developed all-E. *coli* cell-free TXTL system to co-express the array along with FnCas12a and then performed RNA-seq analysis on the purified small RNAs (Marshall et al., 2018). Sequencing showed that the transcribed array was processed into ~44-nt gRNAs similar to prior work (Fonfara et al., 2016; Zetsche et al., 2015), resulting in loss of the 4 nt junctions in most of the sequenced gRNAs. We also observed that the abundance of each gRNA varied widely, potentially impacting the ability of these gRNAs to direct FnCas12a-mediated DNA targeting. Surprisingly, we also observed that the terminal 3’ repeat gave rise to an abundant yet unintended gRNA. Aside from the potential technological complications of creating an errant gRNA, this phenomenon could lead to unintended targeting by natural CRISPR-Cas systems. As a result, natural terminal repeats in Cas12a arrays may have been under selective pressure to accumulate mutations that prevent processing, helping explain the high frequency of mutations in terminal repeats but no other repeats. Correspondingly, analysis of diverse terminal repeats from 14 Type V-A CRISPR-Cas systems revealed that 79% (11/14) of the native terminal repeats harbored mutations known to disrupt Cas12a recognition (**Fig. S1**) (Fonfara et al., 2016; Zetsche et al., 2015). It remains to be seen whether the terminal repeat for the other three CRISPR-Cas systems gives rise to a functional gRNA. These insights aside, we conclude that the generated Cas12a array can be transcribed and undergo processing to form multiple gRNAs.

### CRATES revealed context-dependent loss of FnCas12a spacer activity

We next investigated the targeting activity of the assembled arrays based on plasmid clearance in *E. coli* (**Fig. 2A**). As part of the assay, a targeting or no-spacer CRISPR array plasmid was transformed into cells harboring the FnCas12a plasmid and a plasmid containing the target sequence. Successful plasmid clearance resulted in a large reduction in the number of antibiotic-resistant colonies for the targeting array versus the no-spacer array. We designed three spacers against three distinct target sequences: the *lacZ* promoter (blue), a 5’ portion of the *gfp* coding region (purple), and a 3’ portion of the *gfp* coding region (green). As expected, the CRISPR plasmid with each single spacer cleared only its cognate target plasmid (**Fig. 2A**). The clearance activity was similar in the presence or absence of the junction sequence across all three spacers (**Fig. S2**), ruling out an obvious effect of the junction. We then generated all six configurations of the three-spacer arrays (**Fig. 2A**). Five out of the six arrays cleared all three plasmids. However, one array (PcF-2/3/1) was unable to clear the plasmid with the *lacZ* promoter sequence (blue). Given that the associated spacer drove robust plasmid clearance in the same position in another array (PcF-3/2/1), we attribute the lost activity to the specific context of the spacer in the array.

**Figure 2.**
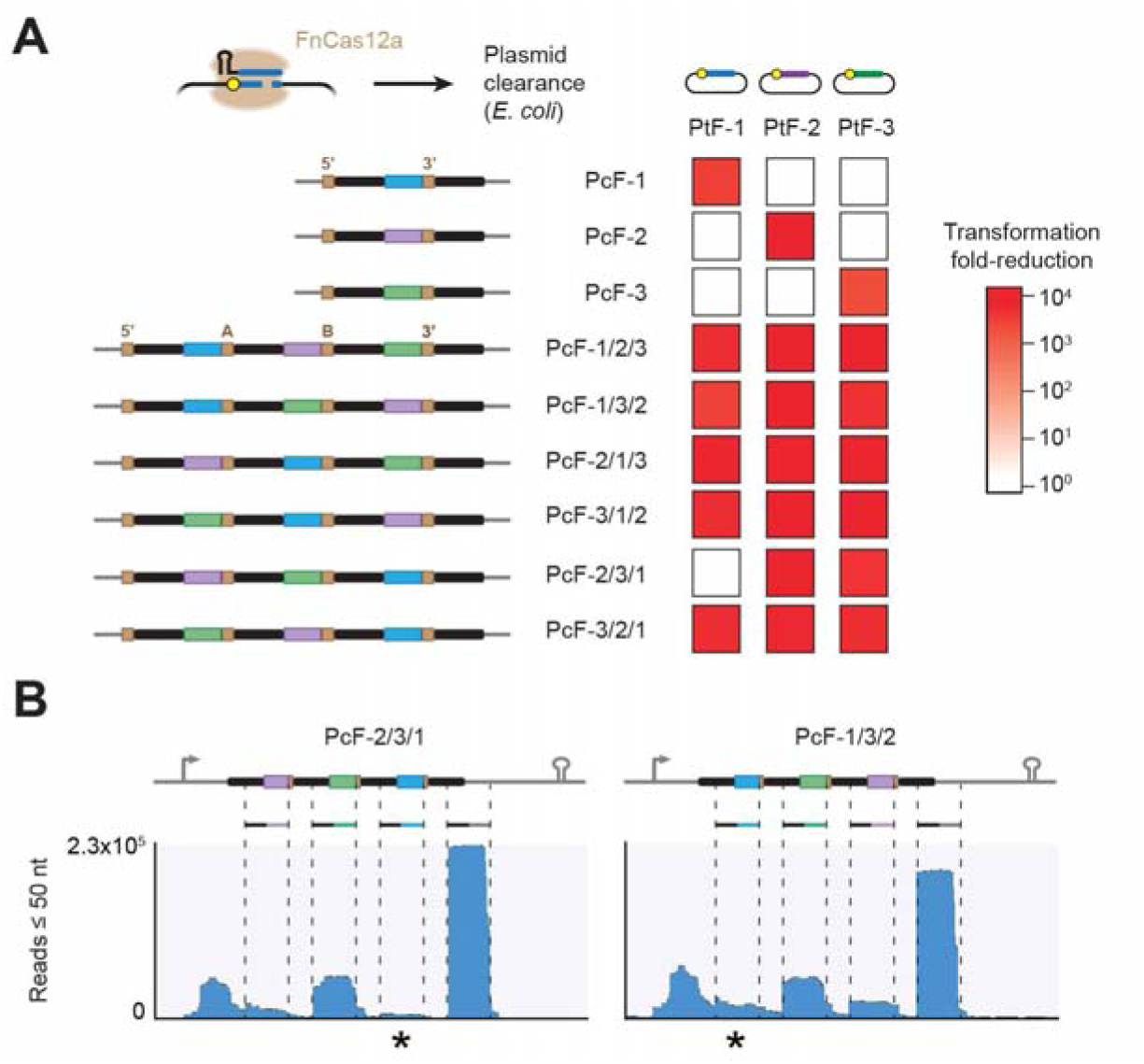
Plasmid clearance with assembled CRISPR-Cas12a arrays reveals context-dependent spacer activity. (A) Multiplexed plasmid clearance by FnCas12a in *E. coli.* Spacers (colored bars) were designed to target a distinct protospacer flanked by a PAM (yellow circle) in a transformed plasmid. *E. coli* cells harboring the FnCas12a plasmid and the target plasmid were transformed with a plasmid encoding the indicated CRISPR array or a no-spacer array, and the fold-change in the number of transformants is reported as a heat map. Sequences of the spacers and junctions in the assembled arrays are shown in Table S3. Values represent the average of at least three independent transformation experiments starting from separate colonies. (B) RNA-seq analysis of three-spacer arrays with differing activity of spacer S1. The assembled array and FnCas12a were expressed in an all-E. *coli* cell-free transcription-translation reactions. Reads of no more than 50 nts were mapped to the original array expression construct. Stars indicate mapped reads for spacer 1. Related to Figures S2-S3 and Tables S2-S3.

RNA-seq analysis of the seven-spacer array revealed widely varying abundances of the processed gRNAs, where a low-abundance gRNA may explain the observed lack of plasmid clearance activity. To evaluate this possibility directly, we performed RNA-seq analysis on the PcF-2/3/1 and PcF-1/3/2 arrays that negligibly or efficiently cleared the plasmid with the *lacZ* promoter sequence, respectively (**Fig. 2B**). The *lacZ* promoter-targeting gRNA in PcF-2/3/1 had the lowest abundance across the two arrays and was ~4-fold lower than the same gRNA in PcF-1/3/2 relative to the middle spacer in each array--all in line with poor plasmid clearance activity. We also noticed that the most-abundant gRNAs were derived from the terminal repeat. These gRNAs could be titrating out available FnCas12a protein, resulting in negligible DNA targeting by less-abundant gRNAs. Given that the terminal repeat is not necessary to generate a functional gRNA (Zetsche et al., 2015), we removed this repeat from both arrays and repeated the plasmid-clearance assay. However, removing the terminal repeat did not appreciably improve plasmid clearance (**Fig. S3A**). Therefore, other factors likely account for the context-dependent loss of targeting activity.

We also explored whether array assembly and multiplexed plasmid clearance could extend to other Cas12a proteins. We chose the Cas12a nuclease from *Acidaminococcus* sp. BV3L6, one of the most commonly used variants of Cas12a (Tang et al., 2017; Zetsche et al., 2015). As expected, we were able to readily generate three-spacer arrays and achieve multiplexed plasmid clearance. All three spacers led to robust plasmid clearance whether in one-spacer arrays or in a three-spacer array (**Fig. S3B**), confirming that CRATES can be extended to different Cas12a nucleases.

### CRATES facilitated multiplexed gene regulation by dCas12a

Following our demonstration of multiplexed DNA cleavage with Cas12a, we next explored the use of CRATES to enact multiplexed gene regulation. Prior work identified three RuvC endonucleolytic domains within FnCas12a, where single point mutations completely (D917A, E1006A) or partially (D1255A) disrupted DNA cleavage activity while retaining DNA binding activity *in vitro* (Fonfara et al., 2016; Zetsche et al., 2015). Single or double point mutations of these domains have been used for programmable gene regulation with Cas12a in bacteria, mammalian cells, and plants (Leenay et al., 2016; Tak et al., 2017; Tang et al., 2017), yet the mutations remain to be compared systematically. We therefore generated variants of Cas12a containing single, double, or triple mutations and then measured DNA targeting of each catalytically-dead variant of FnCas12a (dFnCas12a) based on plasmid clearance or gene repression in *E. coli* (**Fig. 3A**). For either assay, each variant was co-expressed with a single-spacer array targeting the *lacZ* promoter controlling *gfp* on a transformed plasmid. The transformations were performed in an *E. coli* strain lacking this promoter to prevent incidental genome targeting. We found that all tested mutations eliminated any measurable plasmid clearance activity and yielded GFP repression by flow cytometry analysis. Interestingly, one of the single mutations (E1006A) exhibited greatly reduced repression activity that was reversed with additional mutations. Aside from the E1006A single mutation, all other mutations yielded similar strengths of gene repression.

**Figure 3.**
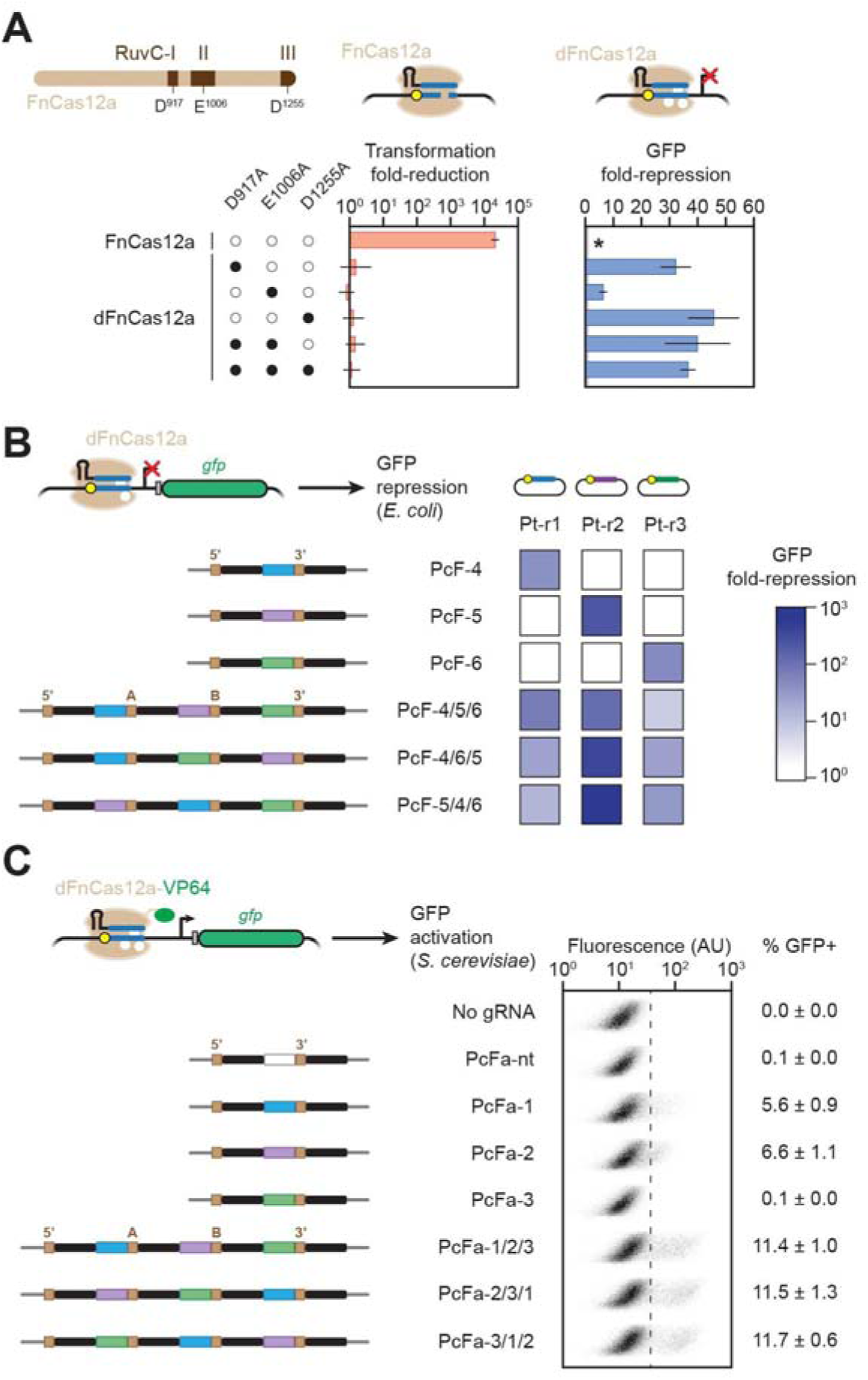
Assembled CRISPR-Cas12a arrays allow multiplexed gene regulation in *E. coli* and yeast. (A) Impact of RuvC mutations on plasmid clearance and gene repression by FnCas12a in *E. coli.* Transformations were conducted as described in Figure 2A. GFP repression was measured by flow cytometry analysis with cells harboring the dFnCas12a plasmid, a plasmid harboring a targeting or no-spacer array, and the GFP reporter plasmid. Values represent the average and S.E.M. of at least three independent experiments from separate colonies. The WT FnCas12a was not tested in the GFP-repression assay (starred) because of its strong plasmid-clearance activity. (B) Multiplexed gene repression with FnCas12a in *E. coli.* See (A) for details, where the D917A, E1006A mutant of FnCas12a was used. Values represent the average and S.E.M. of at least three independent experiments starting from separate colonies. (C) Multiplexed gene activation with a FnCas12a-VP64 fusion in S. *cerevisiae.* Fluorescence distributions were generated from flow cytometry analysis and plotting fluorescence versus side scatter. Values represent the average and S.E.M. of at least three independent experiments starting from separate colonies. All arrays were assembled using the junctions specified in Table S3. Related to Figure S4 and Tables S2-S3.

We selected the triple mutant (D917A, E1006A, D1255A) to evaluate multiplexed gene repression in *E. coli.* We designed spacers to target the *lacZ, lacIq*, or *araB* promoter controlling *gfp* in separate plasmids. We then constructed single-spacer or three-spacer arrays and performed flow cytometry analysis to determine the extent of gene repression for each combination of promoter-arrays in comparison to the no-spacer array (**Fig. 3B**). As expected, each single-spacer array silenced GFP expression only from the cognate promoter, whereas all three-spacer arrays that we tested silenced GFP for all promoters. We also observed some variability in GFP silencing between the tested three-spacer arrays, suggesting some impact of the spacer context, although there was no complete loss of silencing activity.

One of the promising uses of Cas12a is multiplexed gene activation in eukaryotic cells, yet only one study has demonstrated this capability to-date (Tak et al., 2017). We therefore sought to evaluate gene activation in the budding yeast *Saccharomyces cerevisiae* using a fusion between the FnCas12a double mutant (D917A, E1006A) and the transcriptional activator domain VP64. For these experiments, we constructed a new base CRISPR plasmid with the SNR52 snoRNA promoter and a polyT terminator. We then constructed single-spacer or three-spacer arrays targeting different locations around the CYC1 promoter, which drives expression of a downstream chromosomal copy of yeGFP. Flow cytometry analysis showed that each single-spacer array yielded up to 6.6% yEGFP-expressing cells, while the three-spacer arrays consistently yielded 11% yEGFP-expressing cells (**Figs. 3C**, S4). Furthermore, the average fluorescence values were higher for cells with the three-spacer arrays than any of the single-spacer arrays (**Fig. S4**), mirroring the synergistic impact of recruiting multiple activators to the same promoter as observed for dCas9 (Maeder et al., 2013; Perez-Pinera et al., 2013). The observed activation of yEGFP expression was lost when using dFnCas12a without the fused transcriptional activator domain (**Fig. S4**). In total, we showed that CRATES can be used to generate Cas12a arrays for multiplexed gene repression in *E. coli* and multiplexed gene activation in yeast, creating the potential of implementing these arrays in different eukaryotic organisms.

### CRATES is compatible with other Class 2 CRISPR nucleases

We have so far shown that CRATES can be used to construct CRISPR arrays utilized by Cas12a. However, the basis of the assembly method--junctions in the trimmed portion of the spacers--is not limited to Cas12a arrays. Instead, all other Class 2 CRISPR-Cas systems that have been characterized to-date exhibit spacer trimming (Deltcheva et al., 2011; East-Seletsky et al., 2016; Shmakov et al., 2015). This common feature would suggest that CRATES could be compatible with any Class 2 CRISPR nuclease. As a start, we adapted CRATES to assemble CRISPR arrays recognized by the Type II-A single-effector nuclease Cas9 from *Streptococcus pyogenes* (SpCas9), arguably the most widely used CRISPR nuclease to-date (**Fig. 4A**). The major alteration to the base plasmid was placing the repeat upstream of the 5’ restriction site so the assembly junction would fall within 5’ trimmed region of each spacer. Co-expressing the resulting arrays, Cas9, and the tracrRNA in a bacterium with RNase III would lead to processing of the CRISPR array into individual gRNAs (Deltcheva et al., 2011). To demonstrate this functionality, we designed spacers targeting three plasmids respectively. We then cloned three single-spacer arrays and one three-spacer array and performed the plasmid clearance assays in *E. coli.* As expected, the single-spacer arrays only cleared their cognate target plasmid, while the three-spacer assay cleared all three plasmids.

**Figure 4.**
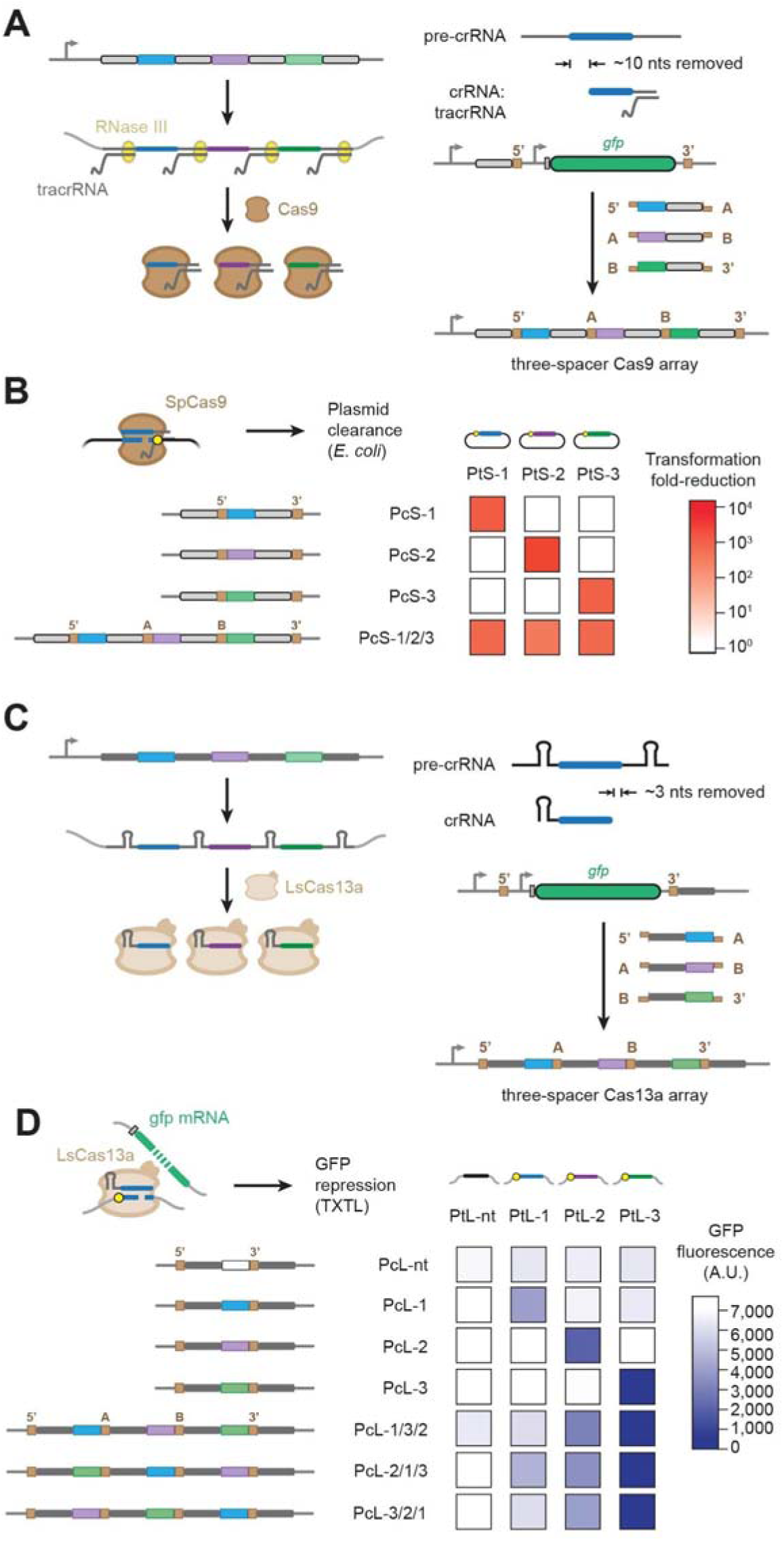
Assembled arrays extend to other Class 2 Cas nucleases. (A) Processing mechanism and assembly scheme for CRISPR-Cas9 arrays. The assembly junction is at the 5’ end of the spacer to match the location of spacer trimming. (B) Multiplexed plasmid clearance by SpCas9 in *E. coli*. See Figure 2A for details. Values represent the average of at least three independent experiments starting from separate colonies. (C) Processing mechanism and assembly scheme for CRISPR-Cas13a arrays. The assembly junction is at the 3’ end of the spacer to match the location of spacer trimming. (D) Multiplexed RNA sensing by LsCas13a in a cell-free transcription-translation system. Reactions were conducted for 16 h following the addition of the LsCas13a plasmid, the indicated array plasmid, a plasmid expressing one of the targets, and the GFP plasmid. Target recognition leads to non-specific degradation of the GFP mRNA by LsCas13a, thereby reducing GFP production. End-point fluorescence measurements are reported. Values represent the average of three TXTL experiments and are representative of at least three experiments conducted on different days. All arrays were assembled using the junctions specified in Table S3. See Figure S5 for the assay and GFP timeccourses. Related to Figure S5 and Tables S2-S3.

As a further demonstration, we used CRATES to explore multiplexed RNA sensing with the Type VI single-effector nuclease Cas13a from *Leptotrichia shahii* (LsCas13a). This nuclease possesses the unique ability to target RNA, where target recognition activates a separate endonuclease domain within LsCas13a that non-specifically degrades cellular RNAs (Abudayyeh et al., 2016). The non-specific activity was recently exploited for the sensitive detection of viral RNAs (Gootenberg et al., 2017, 2018), while this same activity was sufficiently low in human cells to allow programmable and targeted gene silencing with the wild-type nuclease (Abudayyeh et al., 2017). Because the LsCas13a spacer is trimmed on the 3’ end as part of gRNA processing (East-Seletsky et al., 2016; Shmakov et al., 2015), we developed a base assembly construct similar to that for Cas12a (**Fig. 4C**). We then devised an RNA-sensing assay using TXTL, where DNA encoding LsCas13a and the constructed CRISPR array were mixed with DNA expressing a target RNA and a non-targeted GFP reporter (**Fig. S5A,B**). The GFP transcript would undergo non-specific degradation only when the processed gRNA is paired with its target, resulting in a reduction of GFP fluorescence. We created three unique spacers and three complementary targets placed downstream of a constitutive promoter. As expected, GFP expression was reduced compared to the no-spacer array only when the gRNA and its target were both present. Furthermore, the three-spacer arrays reduced GFP in the presence of any of the three target transcripts but not of a non-targeted transcript, resulting in the simultaneous sensing of multiple RNA species (**Figs. 4D, S5C**). Interestingly, some transcripts led to more potent GFP silencing than others, presumably due to ranging secondary structures impacting the accessibility of the target sequence as reported previously (Abudayyeh et al., 2016).

### CRATES-assembled composite arrays mediated coordinated DNA targeting by multiple Cas nucleases

Emerging examples have shown that orthogonal Cas nucleases can be combined to simultaneously perform multiple CRISPR functions, such as conferring immune defense and gene regulation (Esvelt et al., 2013), improving DNA accessibility through binding a proximal site (Chen et al., 2017a), or combining gene disruption and activation in a genome-wide screen (Najm et al., 2018). However, in each case, the gRNAs had to be transcribed from separate expression constructs. We therefore asked if CRATES could be used to generate CRISPR arrays that are processed into gRNAs recognized by multiple Cas nucleases--in what we term “composite” arrays. The simplest arrangement for a composite array would be multiple CRISPR arrays inserted sequentially into the same transcript. Figure 5A illustrates a composite array composed of a two-spacer SpCas9 array followed by a two-spacer FnCas12a array (**Fig. 5B**).

**Figure 5.**
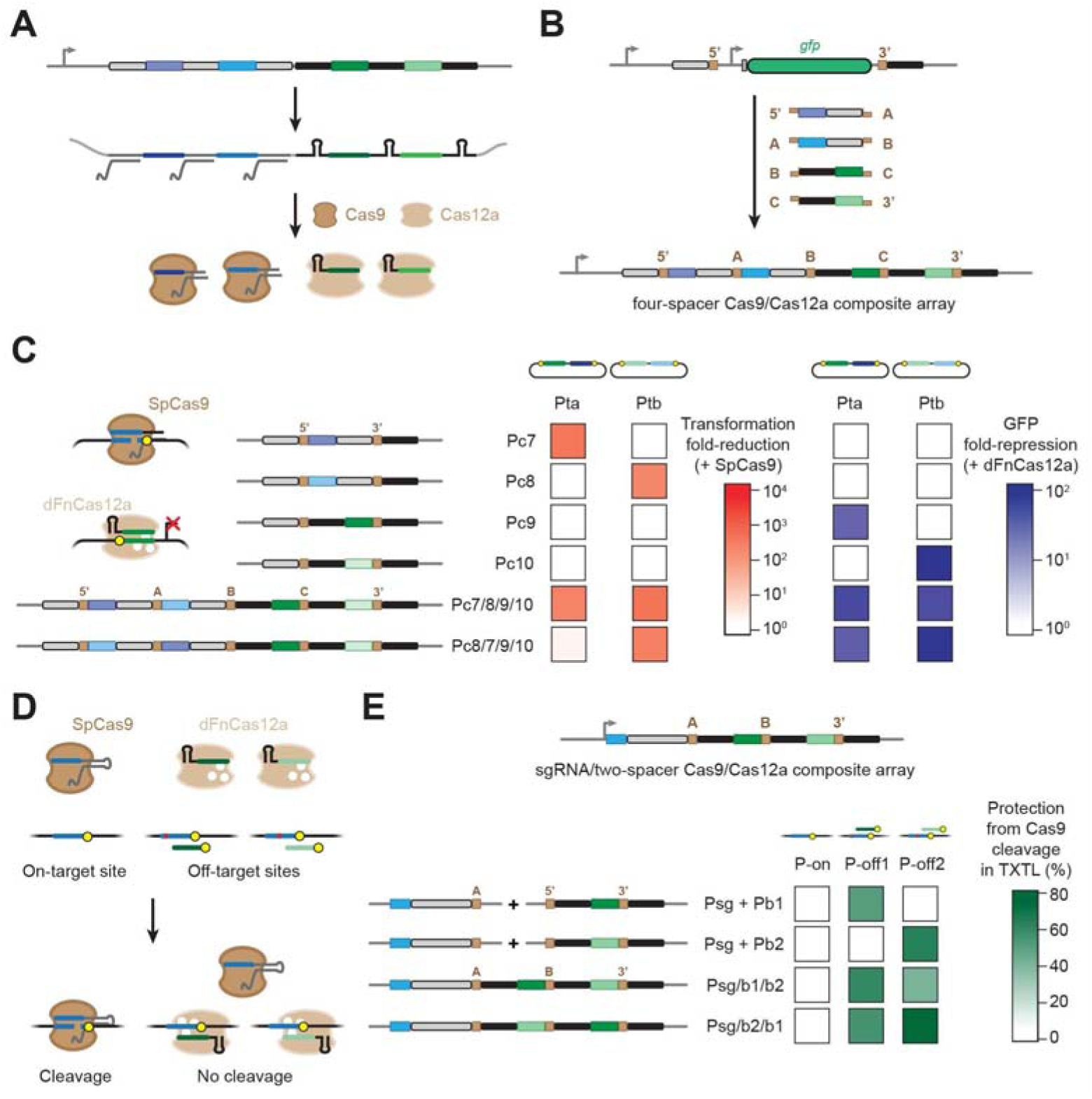
Composite arrays allow recognition and processing by multiple nucleases. (A) Processing of a composite array encoding spacers used by Cas9 and Cas12a. (B) One-step assembly of composite arrays. An array encoding two CRISPR-Cas9 spacers and two CRISPR-Cas12a spacers is shown. The GFP-dropout construct contains an upstream SpCas9 repeat and a downstream FnCas12a repeat to accommodate the orientations of the assembly junctions. (C) Coordinated plasmid clearance by SpCas9 and gene repression by FnCas12a in *E. coli*. To assess plasmid clearance, cells harboring the SpCas9 plasmid and a target plasmid were transformed with a plasmid encoding the indicated CRISPR array or a no-spacer array. To investigate the GFP repression, cells harboring the dFnCas12a (D917A, E1006A, D1255A) plasmid, a target plasmid, and a plasmid encoding the indicated CRISPR array or a no-spacer array were assessed by flow cytometry analysis. Values represent the average of at least three independent experiments starting from separate colonies. See Table S2 and S3 for more information on the target constructs. (D) Enhancing the specificity of DNA cleavage by SpCas9 by blocking off-target sites with dCas12a. Binding of a known off-target location would block Cas9 from accessing this site, thereby reducing unintended cleavage at this site. (E) Enhancing DNA cleavage specificity in a model cell-free TXTL system using a composite array composed of one SpCas9 sgRNA and a two-spacer FnCpf1 array. Protection from cleavage was calculated based on the relative rates of GFP production compared to target and non-targeting controls. See methods for details. Values represent the average of at least three independent TXTL experiments conducted on separate days. See Figure S6 for details of the target sites and a representative set of GFP time-course measurements. Related to Figure S6 and Tables S2-S3.

To explore the technological potential of composite arrays, we first assessed the ability of these arrays to coordinate plasmid clearance by SpCas9 and gene repression by the FnCas12a triple mutant (D917A, E1006A, D1255A) in *E. coli* (**Fig. 5C**). As part of the assessment, we designed two spacers for each nuclease targeting the *lacZ* or *lacIq* promoter controlling *gfp* in the reporter plasmid. We then constructed arrays containing each spacer or all four spacers in two configurations (**Fig. 5C**). *E. coli* cells harboring the SpCas9 or FnCas12a triple-mutant plasmid and either GFP reporter plasmid were transformed with each CRISPR plasmid. We then assessed the transformation efficiency (SpCas9) or GFP fluorescence (dFnCas12a) in comparison to a no-spacer array. As expected, the single spacers yielded plasmid clearance or GFP repression when matched with their nuclease and targeted plasmid, while one of the four-spacer composite arrays (Pc7/8/9/10) yielded plasmid clearance or GFP repression comparable to that of all single-spacer arrays. Interestingly, the other four-spacer composite array with swapped SpCas9 spacers (Pc8/7/9/10) exhibited greatly reduced plasmid clearance of one target plasmid by SpCas9, paralleling the diminished activity of one of the FnCas12a spacers used for plasmid clearance (**Fig. 2A**).

### Composite arrays can improve on-target specificity through coordinated blocking of off-target sites

Following the proof-of-principle demonstration of composite arrays, we sought a more practical application that benefits from coordinated DNA binding and cleavage. One unexplored application is reducing off-target activity by using a catalytically-dead nuclease to bind--and therefore block--known off-target sites. This strategy could be used in combination with existing approaches for diminishing off-target editing such as modified gRNAs or improved-specificity variants of Cas9 (Chen et al., 2017b; Fu et al., 2014; Hu et al., 2018; Kleinstiver et al., 2016a; Slaymaker et al., 2016; Yin et al., 2018), and it could directly exploit a growing suite of experimental techniques for the unbiased detection of off-target sites (Tsai and Keith Joung, 2016). Most importantly, it could allow the selection of target sites that otherwise might have been rejected because of known off-target locations.

We chose SpyCas9 for on-target cleavage and the FnCas12a triple mutant to block off-target sites (**Fig. 5C**). We further selected one of the first examples of off-target cleavage by SpCas9 in human cells: an on-target site in WAS CR-4 (P-on), and two off-target sites in STK25 (P-off1) and GNHR2 (P-off2) (**Fig. S6A**) (Fu et al., 2013). The target sites for FnCas12a were chosen so the R-loop would extend through the NGG PAM recognized by SpCas9, thereby presumably preventing any DNA recognition.

We used TXTL as a rapid and dynamic means to assess whether the catalytically-dead FnCas12a could block SpCas9-mediated cleavage of the off-target sites but not the on-target site. As part of the assay, the target sites were inserted ~170 bps upstream of the P70a promoter controlling *gfp.* Cleavage by SpCas9 would result in rapid degradation by RecBCD and loss of GFP production (**Fig. S6B**) (Maxwell et al., 2018). Using CRATES, we assembled a variant of the composite array encoding the targeting SpCas9 sgRNA upstream of a two-spacer FnCas12a array, where both configurations of the FnCas12a spacers (Psg/b1/b2 and Psg/b2/b1) were tested. We then assayed the cleavage activity of SpCas9 on each site by first adding to the TXTL reaction the FnCas12a triple-mutant plasmid, a reporter plasmid, and either the plasmid encoding a composite array or two plasmids separately encoding the sgRNA and a single-spacer FnCas12a array. We then added the plasmid encoding SpCas9 and measured GFP fluorescence over time. The protection efficiency was then calculated based on the rate of GFP production in comparison to that when expressing a no-spacer FnCas12a array and the targeting sgRNA (0% protection) or a non-targeting sgRNA (100% protection). We found that both tested composite arrays inhibited cleavage by SpCas9 at both off-target sites at efficiencies similar to those when expressing the SpCas9 sgRNA and single-spacer FnCas12a arrays separately (**Fig. 5E**). Critically, there was negligible protection for the on-target site. These results therefore offer an proof-of-principle demonstration of using composite arrays to improve on-target specificity by Cas nucleases.

## DISCUSSION

We have devised and validated a technique we term CRATES for the efficient, one-step generation of large CRISPR arrays. The technique relies on assembling multiple repeat-spacer subunits using defined junction sequences within the trimmed portion of the CRISPR spacers. By specifying the sequence and 5’ or 3’ directionality of the overhang that forms the junction, we could minimize unintended pairing between non-adjacent repeat-spacer subunits. We showed that the technique could generate CRISPR arrays harboring up to seven spacers with high efficiency. While the technique may extend to even larger arrays, the cloning efficiency could be further improved by using different junction lengths (e.g. between 1 and 10 nts for Cas9 spacers) and by subjecting the repeat-spacer oligonucleotides to further purification to remove truncated products.

We showed that the assembled arrays could be utilized by three different single-effector nucleases (Cas9, Cas12a, Cas13a) that yielded multiplexed DNA or RNA targeting in bacteria, eukaryotic cells, and cell-free systems. Given the numerous applications for CRISPR technologies that could benefit from multiplexing--from genome editing, epigenetic regulation, and gene drives to nucleic-acid sensing, antimicrobials, and genetic screens--CRATES has the potential to be widely implemented. Two particularly noteworthy uses are for gene drives and genetic screens. Gene drives utilize a CRISPR nuclease transferring itself and any adjacent genetic cargo to the matching locus in a sister chromosome, thereby allowing the rapid spread of the cargo through a wild population (Esvelt et al., 2014). Because this technique relies on homologous recombination, indel formation through non-homologous end joining (NHEJ) would prevent further editing and create a drive-immune member of the population. By targeting multiple sites in the matching locus through a generated array, disruption of the gene drive through indel formation could be greatly reduced or even eliminated due to the vanishingly small probability that all sites undergo repair through NHEJ simultaneously. In the second example, CRISPR technologies have been used for genome-wide screens based on gene disruption, activation, or repression (Shalem et al., 2015). While these screens principally relied on a library of single sgRNAs, there are recent examples where libraries containing sgRNA pairs were used in human cells to identify synthetic-lethal genes as well as gene pairs that drive cancer proliferation or interact with tumor protein p53 (Najm et al., 2018; Wong et al., 2016). In both examples, the sgRNAs had to be cloned sequentially. CRATES therefore could build on these approaches through the one-step assembly of array libraries that extend beyond two targets, allowing the combinatorial screening of large and potentially redundant factors such as virulence factors, small RNAs, or two-component systems.

We also assembled composite arrays that could be utilized by multiple nucleases (**Fig. 5**). These arrays created the opportunity to enact multiple multiplexed functions by CRISPR nucleases from a single transcript. While expressing or delivering more than one nuclease can be cumbersome, there have been a few applications that benefited from multiple nucleases. In one example, the frequency of gene editing was enhanced by directing catalytically-dead SpCas9 to bind proximally to the site targeted by an orthogonal and catalytically active Cas9 or Cas12a (Chen et al., 2017a). In another example cited above, the SpCas9 and *Staphylococcus aureus* (Sa)Cas9 were used to perform combinatorial screens, where SpCas9 created indels while SaCas9 upregulated gene expression (Najm et al., 2018). In both examples, composite arrays could be used to expand the number of targeting gRNAs produced at one time--whether to enhance editing at one or multiple sites or to increase the number of target genes in a combinatorial screen. We also reported another use of composite arrays based on blocking known off-target sites (**Fig. 5D**). While this approach remains to be demonstrated in cells, it suggests another means to reduce off-target effects that would complement existing strategies and resurrect the use of targets with known off-target sites. This feature would be especially important for edits that can only be achieved through a limited number of target sites, such as for reversing single-nucleotide polymorphisms associated with human disease or performing site-specific integrations (Bulik-Sullivan et al., 2015; Eyquem et al., 2017).

Our assembly scheme depended on the insertion of junctions into the trimmed portions of spacers within a CRISPR array. To-date, spacer trimming has been associated with all Class 2 CRISPR-Cas systems (II, V, VI) and with some Class 1 Type III CRISPR-Cas systems (Hale et al., 2009). On the other hand, Type I CRISPR-Cas systems have not been reported to undergo spacer trimming because of the mechanism of ribonucleoprotein complex assembly (Brouns et al., 2008) and because at least one system has been shown to reject mismatches at the PAM-distal end of the spacer (Szczelkun et al., 2014). CRATES could generate CRISPR arrays for Type I systems by using part of the natural spacer sequence as the junction, where the use of 5’ or 3’ overhangs could be optimized to help eliminate potential cross-interactions between non-adjacent repeat-spacer subunits. This potential limitation aside, Class 2 CRISPR-Cas systems and their single-effector nucleases remain the primary source of CRISPR technologies and would benefit directly from CRATES.

By testing a large cohort of assembled CRISPR arrays, we observed two unique features that could impact their technological potential. One unique feature was spacers being inactive in multi-spacer arrays despite being fully active in single-spacer arrays. In particular, we found an FnCas12a spacer and an SpCas9 spacer that lost their ability to elicit targeted plasmid clearance in the respective context of a three-spacer array (**Fig. 2A**) or a four-spacer composite array (**Fig. 5C**). The effect could not be solely explained by the location of the spacer, as swapping the order of the other spacers restored activity. Instead, the lost activity appears to be context-dependent and could involve interactions with adjacent spacers such as through secondary structure formation. While the lost activity was sparingly observed in our work (individual spacers in 2/26 tested arrays) and was not observed in the few instances in which individual spacers in synthetic arrays were assayed (Luo et al., 2016; Zetsche et al., 2016), it nonetheless could represent an important issue particularly for arrays that cannot be fully validated before they are implemented (e.g. combinatorial screens, editing in multicellular organisms).

Although the exact mechanism underlying the loss of spacer activity remains unclear, we provided evidence linking spacer activity to the low abundance of the resulting gRNA (**Fig. 2B**). Interestingly, natural CRISPR arrays exhibit widely varying abundances of the processed gRNAs that is only partially explained by the proximity to the transcriptional start site (Carte et al., 2014; Deltcheva et al., 2011; Plagens et al., 2014; Zetsche et al., 2015). Instead, gRNA abundance varied widely and was not limited to any single type of CRISPR-Cas system, suggesting that this is a common phenomenon impacting the processing of natural and synthetic gRNAs. More experiments will be necessary to determine why some gRNAs are lower in abundance than others, the connection between abundance and activity, and whether design rules can be elucidated to ensure consistent and high gRNA abundance even for large CRISPR arrays.

The second unique feature that we observed was FnCas12a deriving a gRNA from the terminal repeat in a CRISPR array. This observation was unexpected given that only spacers flanked by two repeats would be expected to become gRNAs. The more concerning ramification is that the terminal repeat-derived gRNA could lead to unintended targeting. From a natural perspective, we provided evidence that terminal repeats for Cas12a arrays have been under negative selection to prevent formation of a gRNA, resulting in mutations within the terminal repeat that disrupt processing. Correspondingly, the one native Cas12a array subjected to RNA-seq analysis--from FnCas12a--did not show any gRNAs derived from the terminal repeat (Zetsche et al., 2015). From a technological perspective, Cas12a gRNAs can be functionally expressed without a 3’ repeat (Kleinstiver et al., 2016b; Zetsche et al., 2015); however, the terminal repeat was important when deriving gRNAs from eukaryotic mRNAs (Zhong et al., 2017). Our results suggest that naturally occurring terminal repeats could be readily used to prevent the generation of an errant gRNA while preserving proper processing and activity of upstream gRNAs.

## ACKNOWLEDGEMENTS

We thank Jörg Vogel and Lars Barquist for critical comments. The pCas9 plasmid was a gift from Luciano Marraffini (Addgene plasmid # 42876). The work was supported through funding from the NIH (1R35GM119561 to C.L.B. and 1DP1DA044359 to A.J.K.), the North Carolina State University Summer Undergraduate Research Grant (to T.D.C.), Agilent Technologies (Gift #3926 to C.L.B.), and the Camille & Henry Dreyfus Foundation (2017-137 to C.L.B.).

## AUTHOR CONTRIBUTIONS

C.L., and R.T.L., and C.L.B. conceived this study. C.L. and C.L.B. designed the experiments. C.L., F.T., R.A.S., and S.R.D. performed the array cloning and experiments with bacteria and TXTL; C.L. and C.L.B. analyzed the data. A.J.K. and T.D.C. conducted the experiments with yeast. R.T.L. analyzed the RNA sequencing data. C.L. and C.L.B. wrote the manuscript, which was read and approved by all authors.

## STAR METHODS

### KEY RESOURCES TABLE

2 KEY RESOURCES TABLE
| REAGENT or RESOURCE | SOURCE | IDENTIFIER |
| --- | --- | --- |
| **Bacterial strains** |  |  |
| *E. coli* BW25113 ΔCRISPR ΔLacOperon | Table S2 | Table S2 |
| *S. cerevisiae* strain YPH500 | Table S2 | Table S2 |
| *E. coli* TOP10 | Table S2 | Table S2 |
| *E. coli* DH5α | Table S2 | Table S2 |
| *E. coli* Novablue | Table S2 | Table S2 |
| *E. coli* Tg1 | Table S2 | Table S2 |
| **Chemicals, Peptides, and Recombinant Proteins** |  |  |
| Q5 DNA Hot Start High-Fidelity DNA Polymerase | New England Biolabs | M0493 |
| BsmBI | New England Biolabs | R0580S |
| BsaI-HF | New England Biolabs | R3535S |
| **Critical Commercial Assays** |  |  |
| myTXTL | Arbor Biosciences | 507024 |
| ZymoPURE Plasmid Midi Prep Kit | Zymo Research | D4200 |
| Gibson Assembly Cloning Kit | New England Biolabs | E5510S |
| NEBuilder HiFi DNA Assembly Cloning Kit | New England Biolabs | E5520S |
| Q5 Site-Directed Mutagenesis Kit | New England Biolabs | E0554S |
| Direct-zol RNA MiniPrep Plus w/ TRI Reagent | Zymo Research | R2071 |
| TURBO DNA-free Kit | Invitrogen | AM1907 |
| T4 Polynucleotide Kinase | New England Biolabs | M0201S |
| RNA Clean and Concentrator Kit | Zymo Research | R1015 |
| Ribo-Zero rRNA Removal Kit (Bacteria) | Illumina | MRZMB126 |
| NEBNext Multiplex Small RNA Library Prep Set for Illumina (Set 1) | New England Biolabs | E7300S |
| **Deposited Data** |  |  |
| RNA-seq analysis of processed FnCas12a arrays | Table S2 | Table S2 |
| **Oligonucleotides** |  |  |
| Primers and cloning oligonucleotides | Table S3 | Table S3 |
| **Recombinant DNA** |  |  |
| Plasmids | Table S2 | Table S2 |
| **Software and Algorithms** |  |  |
| Geneious 10.2.3 | Geneious | N/A |
| **Other** |  |  |
| Detailed protocol for array assembly | Methods S1 | Methods S1 |

### CONTACT FOR REAGENT AND RESOURCE SHARING

Further information and requests should be directed to the Lead Contact, Chase Beisel.

## METHOD DETAILS

### Strains, Plasmids, and Growth Conditions

All strains, plasmids, and oligonucleotides are listed in Table S2 and Table S3. For experiments in *E. coli* and TXTL, SpCas9 was expressed from pCas9 (Addgene #42876) while the other *cas* genes were expressed under the control of the J23108 promoter in a pBAD33 backbone. The targeted plasmids used with the plasmid clearance assays for FnCas12a were constructed by Q5 mutagenesis (New England Biolabs) following the manufacturer’s instructions to remove portions of the pUA66-lacZ plasmid (GFP gene driven by *lacZ* promoter) so that each targeted plasmid only contain one protospacer matching to the designed spacers. The targeted plasmids used with the plasmid clearance assays for SpCas9 were constructed by inserting the fragments containing the PAM and protospacer in between the XhoI and BamHI sites of pUA66. The GFP reporter plasmids used with the gene repression assays were constructed by inserting a constitutive (*lacIQ*) or inducible promoter (*lacZ* or *araB*) into the XhoI and BamHI restriction sites upstream of the *gfp* gene in pUA66 (Zaslaver et al., 2006). The reporter plasmid for *in vitro* RNA detection with LsCas13a was the p70a-deGFP plasmid reported previously (Garamella et al., 2016). The target encoding plasmids were constructed by inserting the protospacer into the plasmid pUA66_PJ23119 by replacing the ORF of GFP gene. Plasmid p70a-deGFP was also used for the off-target binding experiments by inserting each target sequence to upstream of the P70a promoter using Q5 mutagenesis. The Cas12a encoding plasmids were constructed by inserting the open reading frame of FnCpf1 or AsCpf1 into pBAD33 with a constitutive promoter (PJ23108). The plasmids encoding dFnCas12a variants were constructed by Q5 mutagenesis using the wild type FnCas12a plasmid as template. Plasmid used for expressing Cas9 was pCas9 (Addgene # 42876). Plasmid pCas9-notracr was constructed by removing the tracrRNA portion from the plasmid pCas9 using Q5 mutagenesis. The *in vivo* plasmid clearance assays were conducted in CB414, a derivative of *E. coli* BW25113 with the *lacI* promoter through the *lacZ* gene and the endogenous I-E CRISPR-Cas system deleted.

*E. coli* cells were grown in Luria Bertani (LB) medium (10 g/L NaCl, 5 g/L yeast extract, 10 g/L tryptone) at 37°C with shaking at 250 rpm. The antibiotics ampicillin, chloramphenicol, and/or kanamycin were added to maintain any plasmids at 50 μg/mL, 34 μg/mL, and 50 μg/mL, respectively. The inducers Isopropyl β-D-1-thiogalactopyranoside (IPTG) or L-arabinose were added at concentrations of 0.2 mM and 0.2% when specified.

The S. *cerevisiae* strain YPH500 (Stratagene) was used as the background strain for gene activation. Culturing and genetic transformation were done as previously described (Khalil et al., 2012) using either URA3, HIS3, or LEU2 as selectable markers. The reporter plasmid for gene activation was constructed from integrative plasmid pRS406 (Strata-gene) as described previously (Keung et al., 2014). A CEN pRS414 plasmid was used to express the FnCas12a variants from a GPD promoter. To construct FnCas12a-VP64, the FnCas12a double mutant was cloned in between pGPD and VP64 in pRS414 through Gibson assembly. All plasmid constructs were generated using TOP10, Novablue, Tg1, or DH5α electrocompetent cells and verified by PCR and Sanger Sequencing of the inserted sequence.

### Generation of CRISPR arrays

The backbone plasmid used for generating CRISPR arrays for FnCas12a (pFnCpf1GG) was constructed by Gibson assembly to join three PCR fragments together and kill the extra BsmBI site on the scaffold backbone. The three PCR fragments contain a constitutive J23119 promoter, a GFP dropout construct (with promoter and terminator) flanked by two Type IIS BsmBI restriction sites and a direct repeat of FnCas12a, and rrnB terminator, ampicillin resistance gene, and pMB1 origin of replication, respectively. The backbone plasmid used for generating arrays for AsCas12a, and LsCas13a, and the 4-spacer composite arrays were constructed using Q5 mutagenesis to remove the FnCas12a repeat and insert the other corresponding repeats. The backbone plasmid used for generating arrays for SpCas9 was constructed by inserting a PCR fragment with the Cas9 direct repeat and BsmBI sites flanked mRFP dropout construct into Aatll and Hindlll digested pFnCpf1GG. Backbone plasmid for generating composite arrays for blocking off-target cleavage was constructed using Q5 mutagenesis to remove extra nucleotides on the 3’ of the promoter from pFnCpf1GG, so that the transcription starts from the sgRNA of Cas9. Backbone plasmid for generating arrays to use in yeast was constructed by using Gibson Assembly to join three PCR fragments and mutate a BsaI site on the scaffold backbone: BsaI sites flanked GFP dropout construct and two PCR fragments amplified from plasmid pRS413, on which arrays are transcribed from a SNR52 snoRNA promoter. The sequences of the resulting backbone plasmids are available in Table S2. Forward and reverse oligonucleotides encoding one repeat, one spacer, and a 4-nt junction were annealed to form dsDNA with a 5’ and/or 3’ overhangs. 400 fmol of each dsDNA, 20 fmol of backbone plasmid, 1 μl of T7 ligase, and 1 μl of BsmBI or BsaI were added to 2 μl of T4 ligation buffer, then water was added to reach a total volume of 20 μl. A thermocycler was used to perform 25 cycles of digestion and ligation (42 °C for 2 min, 16°C for 5 min) followed by a final digestion step (60°C for 10 min), and a heat inactivation step (80°C for 10 min). The ligation mix was then diluted 1:6 in water and electroporated into competent *E. coli* cells. After transformation and recovery for 1 hour at 37°C with shaking at 250 rpm in SOC media, cells were plated on LB agar containing the appropriate antibiotic and incubated for 16 h. White colonies were then screened for the presence of the correct band size, and the array was validated through Sanger sequencing of the PCR product. See Methods S1 for a detailed protocol and an example of assembling one of the three-spacer FnCas12a arrays.

The no-spacer control used in most of the experiments was generated by inserting a single repeat into the backbone plasmid, resulting in two consecutive repeats with no intervening spacer.

Composite arrays for coordinated plasmid clearance and gene repression were generated using the same method except that the backbone plasmid had a 5’ direct repeat for SpCas9 and and 3’ direct repeat for FnCas12a. Composite arrays for reducing off-targeting were generated by assembling the Cas9 sgRNA and two repeat-spacer dsDNAs into the GFP drop-out backbone. The sgRNA was formed with two annealed oligonucleotides, in line with CRATES.

### *In vivo* plasmid clearance assay

We transformed 50 ng of the plasmid encoding the CRISPR array or a non-targeting control with no spacer into *E. coli* cells harboring a plasmid encoding a Cas protein and another plasmid encoding the gRNA target sequence. After recovering for one hour in SOC at 37°C with shaking at 250 rpm, cells were serially diluted and 10 μl of droplet was plated on LB agar plates with ampicillin, kanamycin, and chloramphenicol. After 16 hours of growth, colony numbers were recorded for analysis.

### *In vivo* gene repression assay

CB414 cells were initially transformed with three compatible plasmids: a plasmid encoding a variant of FnCas12a, a plasmid encoding a CRISPR array or no-spacer control, and a plasmid encoding GFP under the control of the *lacZ, lacIQ*, or *araB* promoter. Overnight cultures of cells harboring the three plasmids were back-diluted to ABS_600_ ~0.01 in LB medium with ampicillin, kanamycin and chloramphenicol and the promoter’s inducer and shaken at 250 rpm at 37 °C until ABS_600_ reach ~0.2. Cultures were then diluted 1:25 in 1X phosphate buffered saline (PBS) and analyzed on an Accuri C6 flow cytometer with C6 sampler plate loader (Becton Dickinson) equipped with CFlow plate sampler, a 488-nm laser, and a 530 +/- 15-nm bandpass filter. GFP fluorescence was measured as described previously (Leenay et al., 2016). Briefly, forward scatter (cut-off of 15,500) and side scatter (cut-off of 600) were used to eliminate non-cellular events. The mean value within FL1-H of 30,0 000 events within a gate set for *E. coli* (Afroz et al., 2014) were used for data analysis.

### Small-RNA library preparation, sequencing, and data analysis

Plasmids encoding FnCas12a and the arrays were added into 9 ul of MyTXTL master mix (Arbor Biosciences) to a final concentration of 5 nM each in a PCR tube and a total volume of 12 ul. Two uM of Chi6 annealed oligos was included in the reaction to prevent any unintentional degradation of the plasmids by RecBCD proteins. The mixture was incubated at 29°C for five hours in a thermocycler, and total RNA was extracted using Direct-zol RNA MiniPrep kit following the manufacturer’s instructions (Zymo Research). After DNase treatment with Turbo DNase (life Technologies), and 3’ dephosphorylation with T4 Polynucleotide Kinase (New England Biolabs), rRNAs were then depleted with the Ribo-Zero Bacteria Kit (Illumina) following the manufacturer’s instructions. RNA libraries were prepared using NEBNext Multiplex Small RNA Library Prep Set for Illumina (New England Biolabs) following the manufacturer’s instructions. Samples were sequenced on a MiSeq machine (Illumina) by the Genomic Science Laboratory at North Carolina State University. The Geneious 10.2.3 software package (Biomatters) was used for data sorting and alignment. Briefly, Fastq reads were trimmed and quality filtered using the BBDuk plugin. Trimmed reads were then aligned to created CRISPR array reference sequences (Fncpf1_7_spacer_array, cF-2/3/1, or cF-1/3/2) using Geneious 10.2.3’ Map to Reference, high sensitivity setting.

### *In vitro* multiplexed RNA detection with Cas13a

Open reading frame of LsCas13a was cloned into pBAD33 with a constitutive promoter. Single or multiple spacer arrays were generated as described. Targeted RNAs were transcribed using a constitutive promoter on the plasmid. A plasmid constitutively express a deGFP protein was used as reporter. The Cas13a encoding plasmid, array encoding plasmid, and target encoding plasmid were added into 9 ul of *MyTXTL* master mix (Arbor Biosciences) to a final concentration of 2 nM, 1 nM and 0.5 nM respectively, to a total volume of 12 ul, and incubated at 37 °C for 2 hours. Then the reporter plasmid was added to a final concentration of 0.5 nM. Aliquots of 5 ul were placed to the wells of 96-well V-bottom plate (Corning Costar 3357) and incubated at 37 °C for 16 hours in a Synergy H1MF microplate reader (BioTek) with kinetic reading every 3 minutes (excitation, emission: 485 nm, 528 nm; gain: 60; lightsource: Xenon Flash).

### Gene activation in *Saccharomyces cerevisiae*

Three single yeast colonies for each strain were picked after plasmid transformations and inoculated into 500 ml of SD-media (synthetic dropout media containing 2% glucose with defined amino acid mixtures) in Costar 96-well assay blocks (V-bottom; 2 ml max volume; Fisher Scientific). The cultures were grown at 30 °C with 250 rpm shaking for 24–48 hr. Cultures were then re-inoculated in SD-complete media to an OD600 of 0.05 to 0.1 and grown at 30 °C with 250 rpm shaking for 12 hours. Cells were treated with 10 mg/ml cycloheximide to inhibit protein synthesis and then assayed for yEGFP expression by flow cytometry. 10,000 events were acquired using a MACSQuant VYB flow cytometer with 96-well plate sampler. Events were gated by forward scatter and side scatter and all values obtained were from three isogenic strains. Plots were generated based on side scatter versus FL1 fluorescence.

### *In vitro* assessment of blocking off-target cleavage

Single guides for Cas9 cleavage, single spacer arrays for dFnCpf1 blocking off-target sites, and composite arrays for cleavage and blocking synchronously were generated as described. The protospacers were cloned into upstream of constitutive promoter of GFP gene on a plasmid. Plasmid encoding dFnCpf1, targeting plasmid for Cas9, targeting plasmid for dFnCpf1 or non-targeting control, and targeted plasmid were added into *MyTXTL* master mix to final concentration of 1 nM, 2 nM, and 0.5 nM respectively, and incubated at 29 °C for 4 hours. Then the SpCas9 encoding plasmid (pCas9-notracr) was added to the mixture to a final concentration of 2 nM and incubated at 29°C for 16 hours in a microplate reader with kinetic reading (excitation, emission: 485 nm, 528 nm) every 3 minutes. Non-targeting control was also tested as a control.

## QUANTIFICATION AND STATISTICAL ANALYSIS

### Plasmid transformation in *E. coli*

Fold reduction was calculated as the ratio of colony-forming units (CFU’s) for cells transformed with the no-spacer array plasmid over that for cells transformed with the CRISPR array plasmid.

### Fluorescence measurements in *E. coli*

GFP fold-repression was calculated using mean fluorescence values, subtracting the fluorescence of *E. coli* cells lacking the GFP reporter plasmid, and dividing the resulting fluorescence value for the no-spacer array by that for the tested CRISPR array.

### Fluorescence measurements in yeast

The fraction of GFP-positive cells were calculated by setting a threshold on FL1 so <0.1% of the cells lacking a gRNA fall within the the GFP-positive bin. The reported fraction is the percentage of cells that fall above the threshold within the entire gated population.

### RNA sensing in TXTL

End-point fluorescence measurements are reported as a heat map.

### Blocking off-target cleavage in TXTL

The transcriptional rate was calculated as the slope of each fluorescence signal curve from 2.5 hours to 4.5 hours after the pCas9-notracr plasmid was added. Protection percentage was calculated as the ratio of the transcriptional rate for CRISPR array over that of the non-targeting control.

## DATA AND SOFTWARE AVAILABILITY

Next-generation sequencing data for the RNA-seq analysis of gRNA abundance will be accessible through NCBI Biosample.

